# Truncated RNA-binding protein production by DUX4-induced systemic inhibition of nonsense-mediated RNA decay

**DOI:** 10.1101/2021.06.28.450189

**Authors:** Amy E. Campbell, Michael C. Dyle, Lorenzo Calviello, Tyler Matheny, Kavitha Sudheendran, Michael A. Cortazar, Thomas Forman, Rui Fu, Austin E. Gillen, Marvin H. Caruthers, Stephen N. Floor, Sujatha Jagannathan

**Affiliations:** Department of Biochemistry and Molecular Genetics, University of Colorado Anschutz Medical Campus, Aurora, CO 80045, USA; Human Technopole, Milan, Italy; RNA Bioscience Initiative, University of Colorado Anschutz Medical Campus, Aurora, CO 80045, USA; Department of Biochemistry, University of Colorado Boulder, Boulder, CO 80309, USA; Department of Craniofacial Biology, University of Colorado Anschutz Medical Campus, Aurora, CO 80045, USA; Medical Scientist Training Program, University of Colorado Anschutz Medical Campus, Aurora, CO 80045, USA; Department of Cell and Tissue Biology, University of California, San Francisco, San Francisco, CA 94143, USA; Helen Diller Family Comprehensive Cancer Center, University of California, San Francisco, San Francisco, CA 94143, USA

**Author notes:** These authors contributed equally (Amy E. Campbell, Michael C. Dyle, Lorenzo Calviello).

**Keywords:** NMD, DUX4, FSHD, translation, splicing

## Abstract

Nonsense-mediated RNA decay (NMD) is a surveillance mechanism that degrades both canonical and aberrant transcripts carrying premature translation termination codons. NMD is thought to have evolved to prevent the synthesis of toxic truncated proteins. However, whether global inhibition of NMD results in widespread production of truncated proteins is unknown. A human genetic disease, facioscapulohumeral muscular dystrophy (FSHD) features acute inhibition of NMD upon expression of the disease-causing transcription factor, DUX4. Here, using a cell-based model of FSHD, we show the production of hundreds of truncated proteins from physiological NMD targets. Using ribosome profiling, we map the precise C-terminal end of these aberrant truncated proteins and find that RNA-binding proteins are especially enriched for aberrant truncations. The stabilized NMD isoform of one RNA-binding protein, SRSF3, is robustly translated to produce a stable truncated protein, which can also be detected in FSHD patient-derived myotubes. Notably, ectopic expression of truncated SRSF3 alone confers toxicity and its downregulation is cytoprotective. Our results demonstrate the genome-scale impact of NMD inhibition. This widespread production of potentially deleterious truncated proteins has implications for FSHD biology as well as other genetic diseases where NMD is therapeutically modulated.

## INTRODUCTION

Nonsense-mediated RNA decay (NMD) degrades transcripts containing premature termination codons (PTCs) that arise from nonsense mutations or RNA processing errors. Through this mechanism, NMD prevents the production of potentially toxic truncated proteins [1]. In addition to its role as a quality control mechanism, NMD also serves to regulate the expression of physiological transcripts that mimic NMD substrates. Such transcripts include a cassette exon containing a PTC, upstream open reading frames, or long 3’ untranslated regions. Additionally, intricate auto- and cross-regulatory feedback loops have evolved that utilize NMD to titrate the level of various splicing factors. An excess amount of these splicing factors facilitates the inclusion of a PTC-containing exon which reduces gene expression (“unproductive splicing”) [2, 3]. Due to its dual role as a quality control and gene regulatory mechanism, NMD efficiency is modulated in a variety of physiological contexts including cell stress, differentiation, and development [4–7]. NMD is also therapeutically targeted to allow the production of certain truncated proteins that retain residual function to counter loss-of-function genetic diseases [8]. In both scenarios, it remains an open question whether NMD inhibition has broader deleterious consequences for the cell.

Depletion of proteins involved in NMD, as well as pharmacological inhibition of NMD, has been shown to upregulate thousands of aberrant transcripts that are typically degraded by NMD [1, 9–11]. However, whether such transcripts produce truncated proteins is not known. At an organismal level, NMD inhibition has been shown to be immunogenic [12, 13], hinting at the production of truncated proteins with neoantigenic epitopes, although the identity of such proteins has not been characterized.

In this study, we took advantage of a cellular model of a human genetic disease, facioscapulohumeral muscular dystrophy (FSHD), where NMD is naturally inhibited and therefore we could investigate the molecular and functional consequences of the loss of NMD. FSHD is a prevalent progressive myopathy caused by misexpression of a double homeodomain transcription factor, DUX4, in skeletal muscle [14, 15]. DUX4 is normally expressed during early embryonic development where it activates the first wave of zygotic gene expression [16–18]. However, in individuals with FSHD, DUX4 is reactivated in the muscle and induces apoptotic death leading to skeletal muscle atrophy [19–22]. We have shown that DUX4 misexpression in muscle cells causes rapid and acute inhibition of NMD followed by proteotoxic stress and, eventually, translation inhibition [23, 24].

Here, we asked whether aberrant RNAs stabilized by DUX4-mediated NMD inhibition produce truncated proteins by performing paired RNA-sequencing (RNA-seq) and ribosome profiling (Ribo-seq) at 0, 4, 8, and 14 hours (h) following the expression of DUX4 in MB135-iDUX4 human skeletal muscle myoblasts, a well-characterized cellular model of FSHD [25, 26]. While RNA-seq measures transcript abundance, Ribo-seq measures ribosome density along an mRNA [27]. Thus, Ribo-seq serves as a proxy for active translation and allows precise delineation of translation start and end sites in order to characterize the protein products made from aberrant RNAs. Using Ribo-seq, we found that hundreds of aberrant RNAs, stabilized by DUX4-mediated inhibition of NMD, are actively translated to produce truncated proteins – particularly truncated RNA-binding proteins (RBPs) and splicing factors. We show that one such truncated splicing factor, serine/arginine-rich splicing factor 3 (SRSF3-TR), is expressed in FSHD muscle cell cultures and contributes to DUX4 toxicity. Thus, our findings demonstrate that NMD inhibition results in the widespread production of truncated proteins with deleterious cellular consequences.

## RESULTS

### DUX4 expression allows functional exploration of the consequences of acute NMD inhibition

Misexpression of DUX4 in skeletal muscle cells inhibits NMD and induces cytotoxicity [22, 26, 28]. To identify time points at which to measure transcript- and translation-level changes induced by DUX4 before the onset of overt cytotoxicity, we utilized a well-characterized doxycycline-inducible DUX4 human myoblast line, MB135-iDUX4 [26], harboring a DUX4-responsive mCherry fluorescent reporter (**Figure 1A**). We live imaged these cells every 15 min for 28 h following doxycycline treatment to induce DUX4 (**Figure 1A**, **Video 1**). Expression of the DUX4-responsive mCherry was rapid and nearly synchronous, with fluorescence detection after 2 h. Cytotoxicity was first observed 9 h following DUX4 induction, with most cells dead or dying by 18 h (**Video 1**). Western blot analysis showed that levels of the key NMD factor UPF1 were reduced to 47% after only 2 h of DUX4 induction and continued to decrease (**Figure 1B**). This confirmed our previous observation that NMD inhibition is an early event during DUX4 expression [23]. Given these data, we chose the time points of 4, 8, and 14 h post-DUX4 induction to investigate the consequences of NMD inhibition by DUX4.

**Figure 1.**
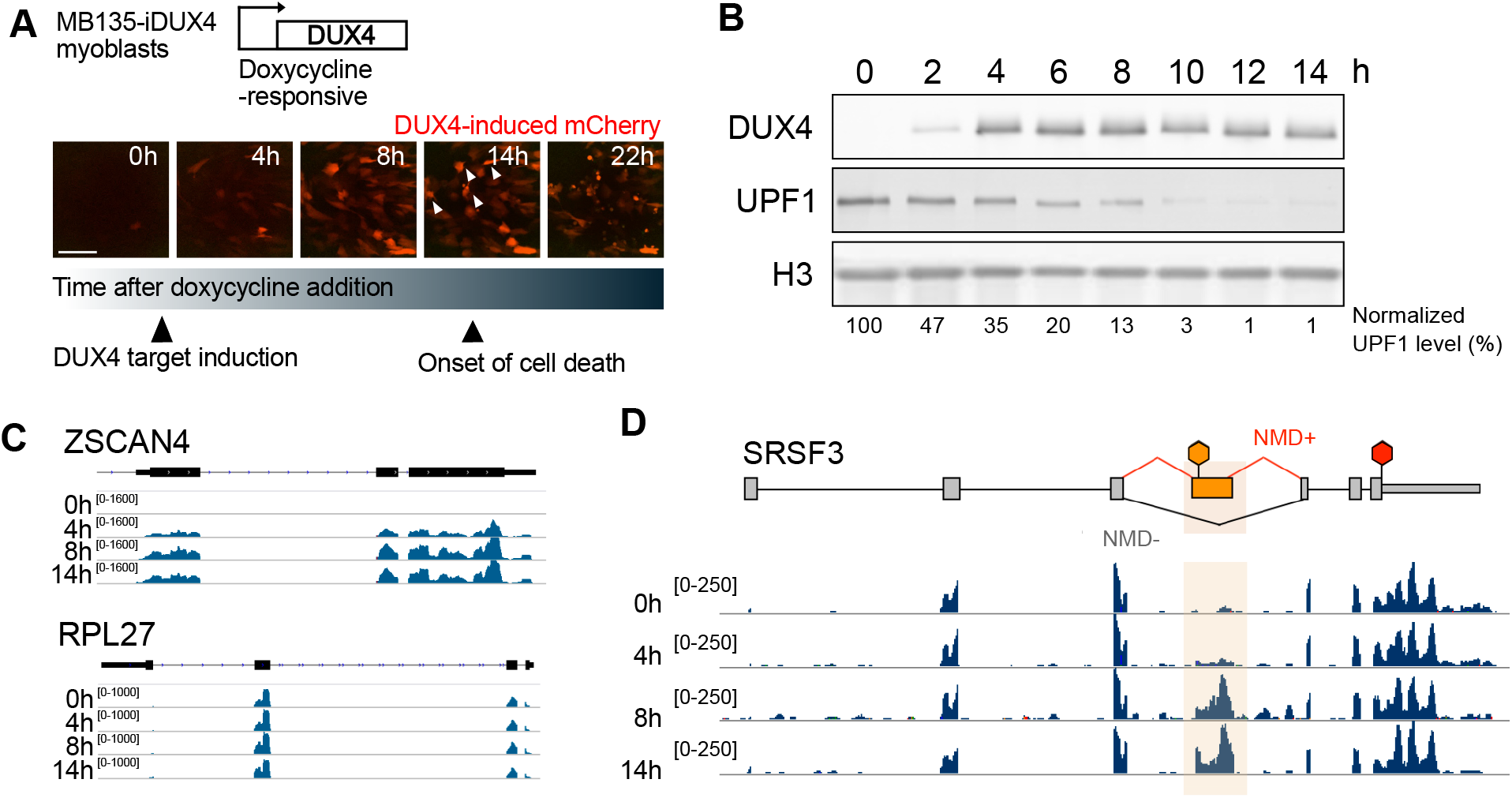
Synchronous expression of DUX4 in MB135-iDUX4 myoblasts enables time course analyses of downstream gene expression changes. (**A**) Representative images from live cell fluorescence microscopy of MB135-iDUX4/ZSCAN4-mCherry myoblasts following treatment with doxycycline to induce DUX4. Arrowheads indicate overtly dying cells. Scale bar, 150 μm. (**B**) Western blot analysis for DUX4, UPF1, and Histone H3 (loading control) over a time course of DUX4 expression following doxycycline induction in MB135-iDUX4 myoblasts. (**C**) RNA-seq read coverage over a time course of DUX4 expression for DUX4 target gene ZSCAN4 (top) and housekeeping gene RPL27 (bottom). (**D**) RNA-seq coverage over SRSF3. The PTC-containing exon 4 is highlighted. The red hexagon indicates the normal stop codon while the orange hexagon denotes the PTC.

First, we examined DUX4-induced transcriptome changes revealed by our RNA-seq dataset (**Supplementary Table 1**). As expected, transcripts of a DUX4 target gene, *ZSCAN4,* were absent in uninduced cells but highly expressed at 4 h and increased with time **(Figure 1C**, top), while the housekeeping gene *RPL27* had constant, robust RNA expression throughout the time course **(Figure 1C**, bottom). Also as expected, aberrant transcript isoforms with PTC-containing exons, such as *SRSF3,* were present at very low levels prior to DUX4 expression but increased in abundance thereafter, appearing as early as 4 h post-induction (**Figure 1D**).

Genome-wide, DUX4 altered the expression of thousands of transcripts, with known DUX4 targets [25] showing increasing upregulation throughout the time course (**Figure S1A**). Using K-means clustering, we grouped the genes significantly altered (defined as absolute log2 fold change > 1 and adjusted p-value < 0.01) at any point during the time course into five clusters (**Figure S1B**) and carried out gene ontology (GO) analysis on each cluster (**Figure S1C**, **Supplementary Table 2**). The genes rapidly induced upon DUX4 expression (Cluster 1) are enriched for negative regulation of cell differentiation, positive regulation of cell proliferation, and DNA-templated transcription, while those rapidly silenced upon DUX4 expression (Cluster 5) are enriched for myogenesis, positive regulation of cell differentiation, and cytoskeleton organization. Together, this is illustrative of a general switch away from a differentiated muscle program and towards a proliferative phenotype, which is consistent with DUX4’s normal role in establishing an early embryonic program. Interestingly, the cluster of genes induced only at the late 14 h time point (Cluster 3) is enriched for GO categories mRNA splicing, ribonucleoprotein transport, ubiquitin-dependent process, unfolded protein response, and hypoxia, which have all been previously reported as major signatures of DUX4-induced gene expression [22, 26, 28]. Together, these RNA-seq data show that the 4, 8, and 14 h time points capture the temporal range of DUX4-induced gene expression changes, and are consistent with early induction of transcriptional responses and late induction of cell stress response.

### Ribosome profiling shows concordance between transcript levels and ribosome occupancy upon DUX4 expression

Previous quantitative analysis of the DUX4-induced proteome via stable isotope labeling by amino acids in cell culture (SILAC) mass spectrometry showed discordant changes at the RNA versus protein level [24] raising the possibility that translation could be modulated upon DUX4 expression. Additionally, at later time points DUX4 induces dsRNA-mediated activation of PKR [29] and stimulates PERK via the unfolded protein response pathway [24], resulting in eIF2α phosphorylation, which is known to inhibit cap-dependent translation [30]. Therefore, we asked whether transcript level changes driven by DUX4 expression were echoed at the level of translation by comparing the RNA-seq and Ribo-seq datasets.

The characteristic 3 nucleotide periodicity exhibited by the ribosome-protected RNA fragments confirmed the high quality of our Ribo-seq data (**Figure S2**). Representative Ribo-seq read coverage plots of the DUX4 target gene, *ZSCAN4,* showed no coverage in uninduced cells, low ribosome density beginning at 4 h, and active translation at 8 and 14 h (**Figure 2A**, top). In contrast, housekeeping gene *RPL27* showed constant, robust translation throughout the time course (**Figure 2A**, bottom). The changes in ribosome-association at these specific genes mirrored the differences seen in their mRNA levels (**Figure 1C**).

**Figure 2.**
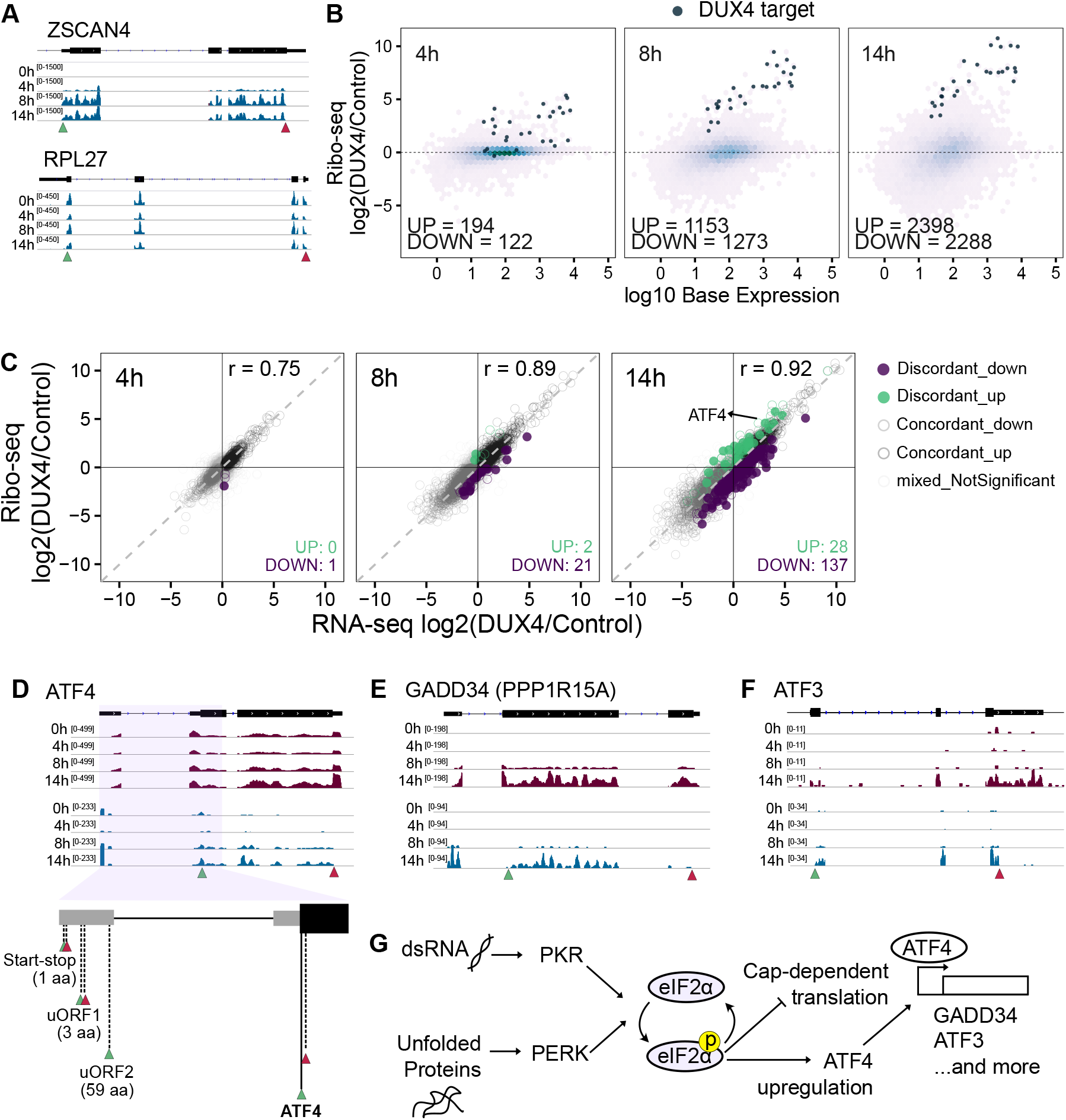
Ribo-seq shows high concordance between transcript levels and translation status. (**A**) Ribo-seq read coverage over a time course of DUX4 expression for DUX4 target gene ZSCAN4 (top) and housekeeping gene RPL27 (bottom). Green triangle, translation start. Red triangle, translation stop. (**B**) M-A plots for Ribo-seq data after 4, 8, and 14 h of DUX4 induction compared to the 0 h control. (**C**) Scatter plot of RNA-seq versus Ribo-seq log2 fold change after 4, 8, and 14 h of DUX4 expression. Significance defined as adjusted p-value < 0.01 for Ribo-seq fold change. (**D**) RNA-seq (top) and Ribo-seq (middle) coverage over ATF4; schematic showing the upstream (uORF) and main open reading frames of ATF4 (bottom). (**E-F**) RNA-seq (top) and Ribo-seq (bottom) coverage over GADD34 (**E**) and ATF3 (**F**). (**G**) A schematic summary of how DUX4 expression influences translation and subsequent cell stress.

On a genome scale, DUX4 altered the translation status for thousands of transcripts, with later time points showing larger differences and known DUX4 targets being translated at increasing levels throughout the time course (**Figure 2B**, **Supplementary Table 3**). Most genes were concordantly up or downregulated at the level of transcript and inferred translation efficiency (**Figure 2C**, Materials and Methods) at all time points with only a small number of genes showing some discordance. GO analysis of the discordantly regulated genes returned significant results only for the gene set that showed a mild translation downregulation at the 14 h time point (n = 137) with pathways such as protein targeting to ER and viral transcription being enriched which, strikingly, were driven entirely by a group of ribosomal protein-encoding genes (**Supplementary Table 3**). This mild downregulation of translation is consistent with induction of the integrated stress response (ISR) pathway and DUX4-mediated eIF2α phosphorylation [24, 29]. A caveat of Ribo-seq to keep in mind here is that it may not capture the true absolute change in translation efficiency induced by ISR, and therefore the translation downregulation we observe could be an underestimate.

Nonetheless, we observe several hallmarks of ISR activation at 14 h, including robust translation upregulation of ATF4 (**Figure 2C-D**). Specifically, the start-stop regulatory element in the 5’ untranslated region of ATF4, which modifies downstream re-initiation to enhance ATF4’s inducibility under stress [31], is highly occupied by ribosomes at 14 h (**Figure 2D**). The resultant upregulation of ATF4 protein can account for the subsequent transcriptional and translational induction of the ATF4 targets GADD34 and ATF3 (**Figure 2E-F**). These data confirm the DUX4-induced phosphorylation of eIF2α via PKR and PERK activation culminate in a block in capdependent translation, and stimulation of robust ISR signaling at the late 14 h time point (**Figure 2G**). Yet, in large part DUX4-induced changes in transcript level are mirrored in their ribosome occupancy.

### DUX4 causes widespread truncated protein production

Having shown that most transcripts induced by DUX4 are robustly translated, and that NMD inhibition is an early consequence of DUX4 expression, we sought to determine whether truncated proteins are produced from PTC-containing RNAs stabilized by DUX4-mediated NMD inhibition. Strikingly, when we compared the RNA-seq and Ribo-seq reads at candidate genes with NMD isoforms, IVNS1ABP, SRSF3, SRSF6, and SRSF7, we saw robust coverage across the PTC-containing exon with reads stopping at the PTC (**Figure 3A**). This clearly indicates the translation of truncated proteins from NMD isoforms.

**Figure 3.**
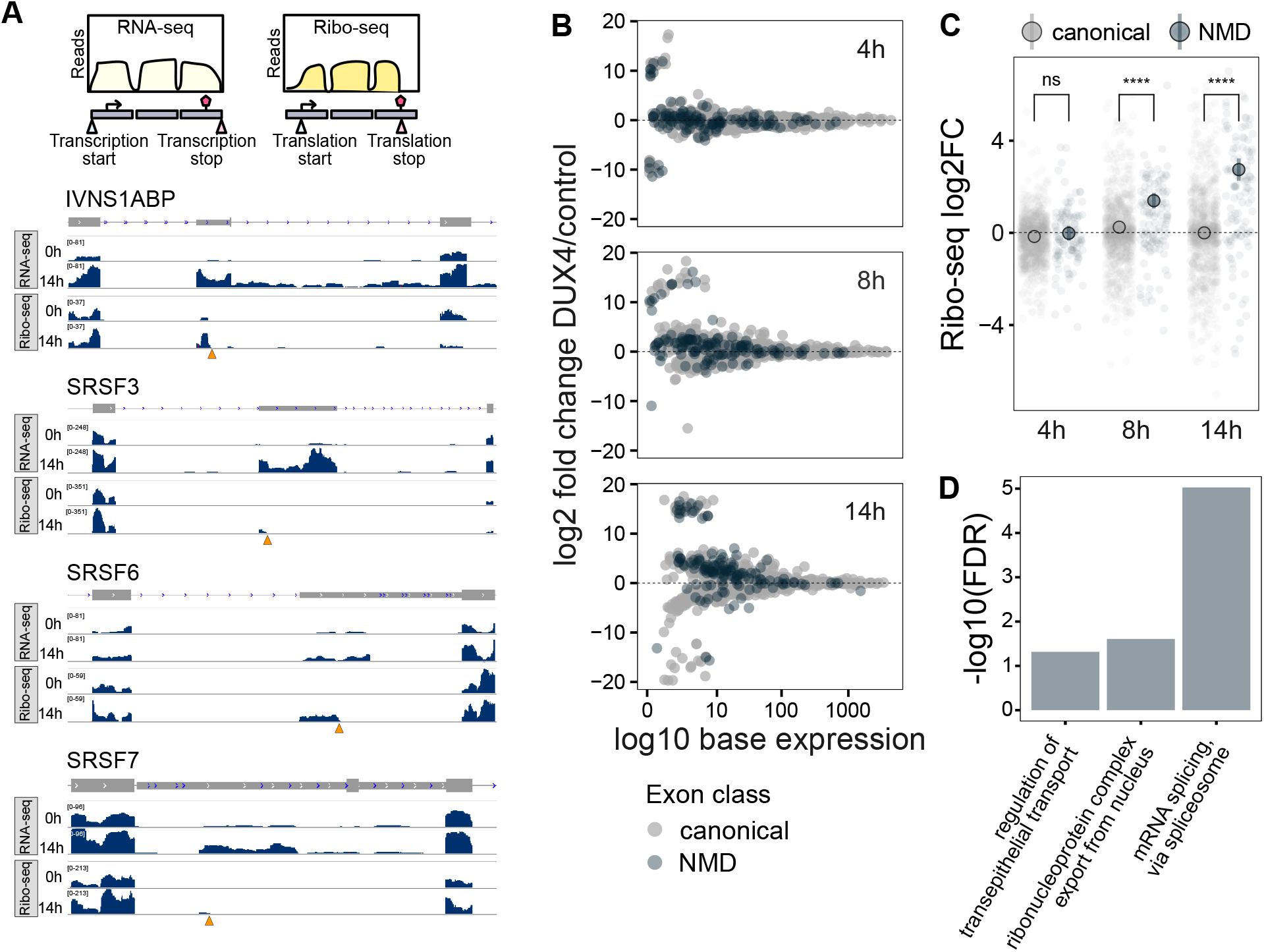
Exon-level analysis of *de novo* identified ORFs from Ribo-seq data demonstrates robust translation of NMD-targeted aberrant RNAs. **(A)** Schematic representation of the paired RNA-seq and Ribo-seq experiment (top) and RNA-seq and Ribo-seq coverage over splicing-related genes IVNS1ABP, SRSF3, SRSF6, and SRSF7 (bottom). Orange triangles denote PTCs. (**B**) M-A plot of exon-level analysis of Ribo-seq data from 4, 8, and 14 h of DUX4 induction. The x-axis represents mean expression calculated at the level of each exon within a gene. (**C**) Exonlevel log2 fold change Ribo-seq values at 4, 8, or 14 h of DUX4 expression for canonical and NMD exons. Statistical testing was performed using a two-sided Wilcoxon test. (**D**) GO analysis results of selected gene sets (biological process complete) for all NMD targets that are translated at 14 h.

To systematically ask if and when aberrant RNAs are translated on a genome-level, we used ORFquant [32], a new pipeline that identifies isoform-specific translation events from Ribo-seq data. We then used DEXSeq [33] to conduct exon-level differential analysis on the set of ORFquant-derived open reading frames, using Ribo-seq data. This analysis identifies changes in relative exon usage to measure differences in the expression of individual exons that are not simply the consequence of changes in overall transcript level. After 4 h of DUX4 induction 397 genes showed differential expression of specific exons, of which 24 are predicted NMD targets (**Figure 3B**), whereas later time points showed a greater number of exons (n = 96 at 14 h) that are unique to NMD targets as differentially expressed (**Supplementary Table 4**). We grouped exons based on their NMD target status and calculated their fold change in ribosome footprints at 4, 8, and 14 h of DUX4 expression compared to the 0 h time point (**Figure 3C**). We observed a progressive and significant increase in the translation status of NMD-targeted exons, but not canonical exons, at 8 and 14 h, confirming the translation of stabilized aberrant RNAs in DUX4-expressing cells.

Translation of an NMD target typically generates a prematurely truncated protein. To ask how the specific truncated proteins being produced in DUX4-expressing myoblasts might functionally impact cell homeostasis, we conducted GO analysis of the 74 truncated proteins being actively translated at 14 h of DUX4 induction (**Figure 3D**, **Supplementary Table 4**). Strikingly, the truncated proteins are enriched for genes encoding RBPs involved in mRNA metabolism and specifically, splicing (**Supplementary Table 4**). Thus, not only are NMD targets stabilized by DUX4 expression, but they also produce truncated versions of many RBPs, which could have substantial downstream consequences to mRNA processing in DUX4-expressing cells.

### Truncated SRSF3 is present in FSHD myotubes and contributes to cytotoxicity

To explore the role of truncated proteins in DUX4-induced cellular phenotypes, we chose SRSF3 for further characterization. SRSF3 is an SR family protein that possesses an N-terminal RNA-binding RNA recognition motif (RRM) and a C-terminal arginine/serine (RS)-rich domain responsible for protein-protein and protein-RNA interactions. The NMD isoform of SRSF3 encodes a truncated protein (SRSF3-TR) that lacks most of the RS domain and has been previously implicated in a variety of human pathologies [34–38]. Examination of our Ribo-seq data revealed robust expression and translation of SRSF3 NMD-targeted exon 4 that ends at the site of the PTC (**Figure 4A**).

**Figure 4.**
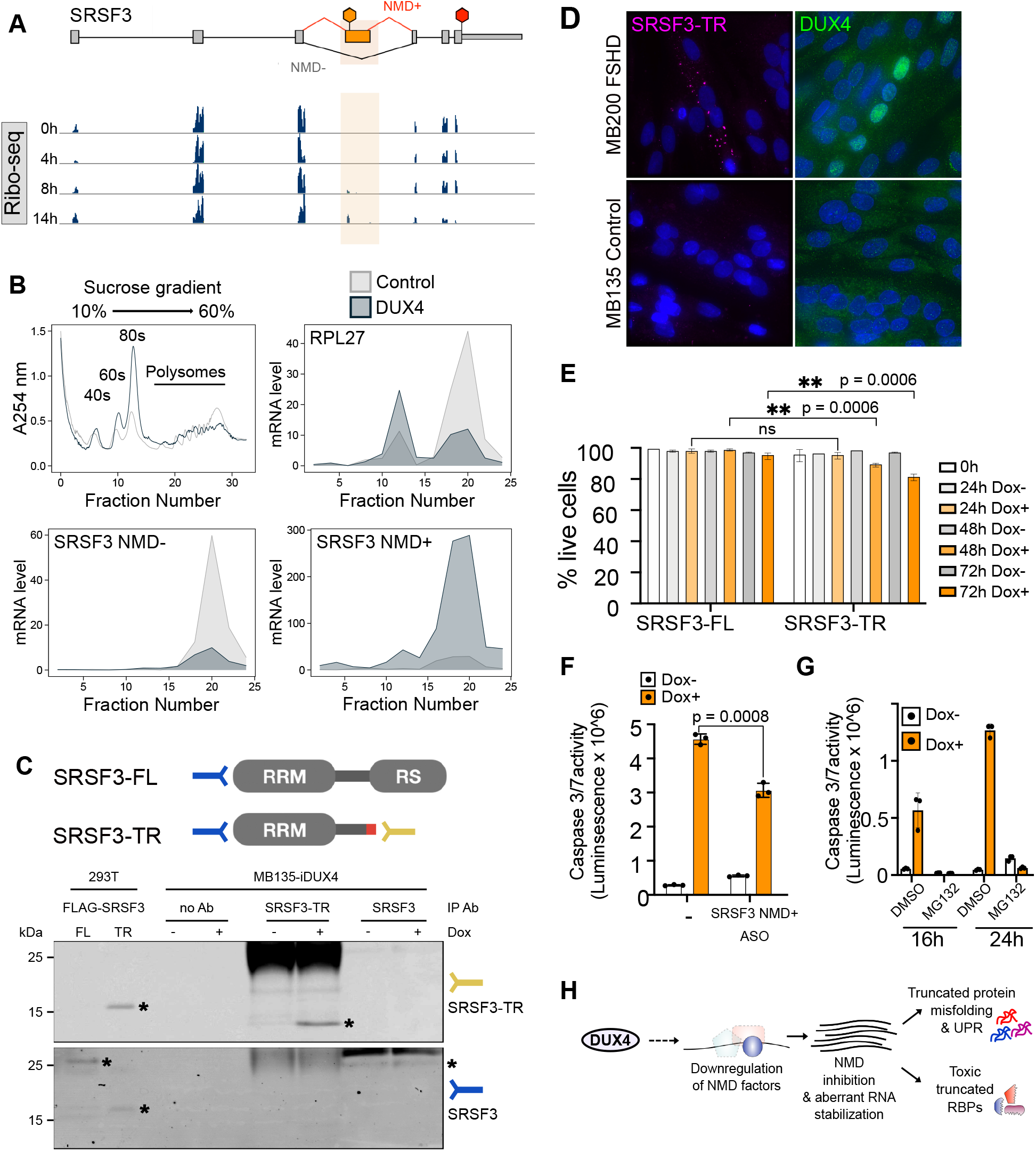
Truncated SRSF3 protein could disrupt RNA processing in FSHD myotubes. (**A**) Ribo-seq coverage over SRSF3. The PTC-containing exon 4 is highlighted. The red hexagon indicates the normal stop codon while the orange hexagon denotes the PTC. (**B**) Absorbance at 254 nm across a sucrose density gradient of lysates from control MB135-iDUX4 myoblasts and MB135-iDUX4 myoblasts expressing DUX4 for 14 h (top left). RT-qPCR measurement of RPL27, SRSF3 NMD-, and SRSF3 NMD+ mRNA levels in DUX4-expressing myoblasts relative to control myoblasts from collected fractions (remaining panels). (**C**) Detection of SRSF3 and SRSF3-TR in whole cell extracts from 293T cells exogenously expressing FLAG-tagged full-length (FL) or truncated (TR) SRSF3 as compared to protein lysates from MB135-iDUX4 myoblasts treated with (+) or without (-) doxycycline (Dox) to induce DUX4 and immunoprecipitated with a custom anti-SRSF3-TR antibody, no antibody (Ab), or a commercial SRSF3 antibody. IP, immunoprecipitation. Asterisks denote proteins of interest. (**D**) Immunofluorescence staining in MB135 control and MB200 FSHD myotubes differentiated for 72 h and stained with DAPI (blue) and rabbit anti-DUX4 (green) or custom rabbit anti-SRSF3-TR (pink) antibody. (**E**) Trypan blue exclusion-based live cell counts in myoblasts expressing doxycycline (Dox)-inducible full-length (FL) or truncated (TR) SRSF3. Error bars denote the standard deviation from the mean of three biological replicates. (**F**) Caspase 3/7 activity following ASO-mediated knockdown of SRSF3 NMD+ in MB135-iDUX4 myoblasts left untreated (Dox-) or treated with doxycycline for 16 h (Dox+) to induce DUX4. Error bars denote the standard deviation from the mean of three biological replicates. (**G**) Cell viability as measured by Caspase 3/7 activity following co-treatment with doxycycline (Dox) to induce DUX4 and proteasome inhibition via MG132 treatment for 16 or 24 h. (**H**) Schematic representation of a working model where DUX4-induced downregulation of NMD factors stabilizes aberrant RNAs producing truncated RBPs and misfolded proteins that trigger the unfolded protein response and toxicity.

To determine the translation status of the aberrant SRSF3 transcript stabilized by DUX4, we carried out polysome profiling using sucrose density gradient separation. The polysome profile after 14 h of DUX4 expression compared to control showed a higher fraction of 80S compared to polysomes (**Figure 4B**, top left). This is consistent with our prior observation of eIF2α phosphorylation [24, 29] and resultant global downregulation of translation at 14 h. We extracted RNA from various fractions and profiled RNA levels of specific transcripts by qPCR. *RPL27* mRNA was predominantly ribosome-bound in control cells but this partially shifted to monosomes in DUX4-expressing cells (**Figure 4B**, top right). The loss of transcripts from polysome fractions was even more stark for the normal SRSF3 isoform (SRSF3 NMD-) which decreased substantially in DUX4-expressing myoblasts compared to control (**Figure 4B**, bottom left). In striking contrast to both RPL27 and SRSF3 NMD-, the NMD-targeted isoform of SRSF3 (SRSF3 NMD+) showed a massive increase in heavy polysomes in DUX4-expressing cells (**Figure 4B**, bottom right). These data show that aberrant SRSF3 mRNA is being actively translated into truncated protein in DUX4-expressing myoblasts and validate the ribosome footprints found on the NMD+ isoform.

To determine if we could stably detect truncated SRSF3 protein in cells, we generated an antibody recognizing a 10 amino acid C-terminal neo-peptide unique to SRSF3-TR. This custom SRSF3-TR antibody was able to recognize FLAG-tagged SRSF3-TR exogenously expressed in 293T cells and endogenous SRSF3-TR immunoprecipitated from DUX4-expressing MB135-iDUX4 myoblasts (**Figure 4C**). We also used a commercial SRSF3 antibody that recognizes an N-terminal epitope common to both the full-length and truncated SRSF3. This antibody detected both exogenously expressed full-length and truncated FLAG-SRSF3, and endogenous full-length SRSF3, but was insufficient to visualize endogenous SRSF3-TR (**Figure 4C**), possibly due to lower affinity for this protein isoform in an immunoprecipitation assay. To determine if SRSF3-TR was present in FSHD myotubes expressing endogenous levels of DUX4, we carried out immunofluorescence for SRSF3-TR or DUX4 in differentiated FSHD and control muscle cells. While there was no detectable SRSF3-TR staining in control cells, in DUX4-expressing FSHD cultures SRSF3-TR appeared in cytoplasmic puncta (**Figure 4D**). Thus, not only is SRSF3 NMD+ isoform translated robustly in DUX4-expressing cells, its protein product, SRSF3-TR can be detected in both DUX4-expressing MB135-iDUX4 myoblasts and in FSHD patient-derived myotubes.

To ask if expression of SRSF3-TR is deleterious to cells, we exogenously expressed FLAG-tagged full-length or truncated SRSF3 in healthy muscle cells via lentiviral transduction. We found that SRSF3-TR, but not SRSF3-FL, reduced the viability of MB135 myoblasts (**Figure 4E**). To specifically knock down the SRSF3-TR isoform in DUX4-expressing myoblasts, we screened several antisense oligonucleotides (ASOs) in order to identify one (a thiomorpholino 2’-deoxyribonucleotide 3’-thiophosphate oligonucleotide chimera [39, 40]) that lowered the SRSF3-TR isoform without significantly impacting DUX4 transcript level or target expression (**Figure S3**). Treatment of DUX4-expressing myoblasts with this thiomorpholino oligonucleotide resulted in a 34% reduction in cell death compared to untreated cells (**Figure 4F**). Finally, we found that blocking the proteasome, which is the primary mediator of NMD inhibition by DUX4, robustly rescues DUX4 toxicity (**Figure 4G**). These results suggest that truncated proteins confer toxicity to muscle cells via a gain-of-function mechanism. Significantly, this mechanism is potentially additive across the different species of truncated proteins that are robustly made upon DUX4 expression in myoblasts (**Figure 4H**).

## DISCUSSION

Loss of NMD leads to the stabilization of aberrant RNAs [41]. However, it is not known whether these aberrant RNAs are translated and what proteins they might produce. Here, we paired RNA-seq and Ribo-seq across a time course of DUX4 expression in human skeletal muscle myoblasts to demonstrate that DUX4-induced NMD inhibition indeed causes truncated protein production at a genome level.

The production of truncated proteins upon NMD inhibition by DUX4 has multiple implications at both molecular and functional levels. Protein truncation could result in a dominant negative function that inhibits the activity of the remaining, cell critical full-length protein. Truncated proteins might misfold and facilitate the formation of protein aggregates. And finally, some truncated proteins contain unique C-terminal extensions, or neo-peptides, that could serve as novel antigens and might induce inflammation. Hence, when NMD is modulated as a therapeutic intervention for genetic diseases, it is important to consider whether truncated proteins are produced as a consequence, and whether this might have a negative impact. In physiological contexts where NMD efficiency is suppressed without negative consequences, cells may possess mechanisms that counter truncated protein production. Further investigation of the suppression or tolerance of truncated proteins in these contexts would reveal new mechanisms that enable the protein quality control rheostat to be adjusted to deal with variable NMD efficiencies.

In FSHD, there is evidence for truncated proteins contributing to myotoxicity via all of the above mechanisms. Here, we show gain-of-function toxicity for SRSF3-TR. Prior work has demonstrated protein aggregation [29, 42, 43], as well as immune cell infiltration [44–47] in FSHD muscle. Our results suggest that these effects could be due to truncated proteins and neoantigenic epitopes. In addition, many of the identified DUX4-induced truncated proteins are RBPs and splicing factors. It is well-established that DUX4 alters RNA splicing [22, 24, 26, 28] and therefore interesting to speculate that truncated RBPs and splicing proteins might be responsible for inducing global RNA processing defects [48]. Such misprocessing would generate aberrant RNAs that could act to further overwhelm the already inhibited NMD pathway.

In summary, we demonstrate the widely-held but previously unproven assumption that NMD inhibition indeed results in the production of truncated proteins with deleterious cellular consequences. In doing so, we provide a framework to interpret the multifaceted phenotypes observed in FSHD as a potential result of NMD inhibition. Our findings provide a critical missing piece in the understanding of this essential quality control mechanism in both disease and physiology, which has implications for treatment of genetic diseases.

## MATERIALS AND METHODS

### RESOURCE AVAILABILITY

#### Lead contact

Further information and requests for resources and reagents should be directed to and will be fulfilled by the lead contact, Sujatha Jagannathan.

#### Materials availability

The cell lines and antibody generated in this study are available upon request. Plasmids generated in this study have been deposited to Addgene (plasmid #171951, #171952, #172345, and #172346).

#### Data and code availability

The RNA-seq and Ribo-seq data generated during this study are available at GEO (accession number GSE178761). The code generated during this study are available at GitHub (https://github.com/sjaganna/2021-campbell_dyle_calviello_et_al).

### EXPERIMENTAL MODEL AND SUBJECT DETAILS

#### Cell lines and culture conditions

293T cells were obtained from ATCC (CRL-3216). MB135, MB135-iDUX4, MB135-iDUX4/ZSCAN4-mCherry, and MB200 immortalized human myoblasts were a gift from Dr. Stephen Tapscott and originated from the Fields Center for FSHD and Neuromuscular Research at the University of Rochester Medical Center. MB135-iDUX4 cells have been described previously [26]. MB135-iFLAG-SRSF3-FL, and MB135-iFLAG-SRSF3-TR immortalized human myoblasts were generated in this study. All parental cell lines were authenticated by karyotype analysis and determined to be free of mycoplasma by PCR screening. 293T cells were maintained in Dulbecco’s Modified Eagle Medium (DMEM) (Thermo Fisher Scientific) supplemented with 10% EqualFETAL (Atlas Biologicals). Myoblasts were maintained in Ham’s F-10 Nutrient Mix (Thermo Fisher Scientific) supplemented with 20% Fetal Bovine Serum (Thermo Fisher Scientific), 10 ng/mL recombinant human basic fibroblast growth factor (Promega), and 1 μM dexamethasone (Sigma-Aldrich). MB135-iDUX4/ZSCAN4-mCherry and MB135-iDUX4 myoblasts were additionally maintained in 2 μg/mL puromycin dihydrochloride (VWR). MB135-iFLAG-SRSF3-FL and -TR myoblasts were additionally maintained in 10 μg/mL blasticidin S HCl (Thermo Fisher Scientific). Induction of DUX4 and SRSF3 transgenes was achieved by culturing cells in 1-2 μg/mL doxycycline hyclate (Sigma-Aldrich). Differentiation of myoblasts into myotubes was achieved by switching the fully confluent myoblast monolayer into DMEM containing 1% horse serum (Thermo Fisher Scientific) and Insulin-Transferrin-Selenium (Thermo Fisher Scientific). All cells were incubated at 37 °C with 5% CO_2_.

### METHOD DETAILS

#### Cloning

pTwist-FLAG-SRSF3_Full.Length_Codon.Optimized and pTwist-FLAG-SRSF3_Truncated_Codon.Optimized plasmids were synthesized by Twist Bioscience. To construct pCW57.1-FLAG-SRSF3_Full.Length_Codon.Optimized-Blast and pCW57.1-FLAG-SRSF3_Truncated_Codon.Optimized-Blast plasmids, the SRSF3 open reading frames were subcloned into pCW57-MCS1-P2A-MCS2 (Blast) (a gift from Adam Karpf, Addgene plasmid #80921) [49] by restriction enzyme digest using EcoRI and BamHI (New England Biolabs).

#### Antibody generation

Purified SRSF3-TR peptide (Cys-PRRRVTIMSLLTTL) was used as an immunogen and polyclonal rabbit anti-SRSF3-TR antibody production was done in collaboration with Pacific Immunology (Ramona, CA). The antisera from all animals were screened for reactivity by ELISA against the immunogen and with western blots and immunofluorescence against transfected SRSF3-TR.

#### Transgenic cell line generation

Lentiviral particles expressing doxycycline-inducible FLAG-SRSF3-FL or -TR transgenes were generated by co-transfecting 293T cells with the appropriate lentivector, pMD2.G (a gift from Didier Trono, Addgene plasmid #12259), and psPAX2 (a gift from Didier Trono, Addgene plasmid #12260) using Lipofectamine 2000 Transfection Reagent (Thermo Fisher Scientific). To generate polyclonal SRSF3 transgenic cell lines, MB135 myoblasts were transduced with lentivirus in the presence of 8 μg/mL polybrene (Sigma-Aldrich) and selected using 10 μg/mL blasticidin S HCl.

#### Plasmid transfections

293T cells were transfected with pTwist-FLAG-SRSF3_Full.Length_Codon.Optimized and pTwist-FLAG-SRSF3_Truncated_Codon.Optimized plasmids using Lipofectamine 2000 Transfection Reagent following the manufacturer’s instructions.

#### Live cell imaging

MB135-iDUX4/ZSCAN4-mCherry myoblasts were induced with doxycycline hyclate to turn on DUX4 expression and subjected to time lapse imaging using the IncuCyte S3 incubator microscope system (Sartorius). Images were collected every 15 min from the time of doxycycline addition (t = 0 h) to 28 h.

#### RNA extraction and RT-qPCR

Total RNA was extracted from whole cells using TRIzol Reagent (Thermo Fisher Scientific) following the manufacturer’s instructions. Isolated RNA was treated with DNase I (Thermo Fisher Scientific) and reverse transcribed to cDNA using SuperScript III reverse transcriptase (Thermo Fisher Scientific) and random hexamers (Thermo Fisher Scientific) according to the manufacturer’s protocol. Quantitative PCR was carried out on a CFX384 Touch Real-Time PCR Detection System (Bio-Rad) using primers specific to each gene of interest and iTaq Universal SYBR Green Supermix (Bio-Rad). The expression levels of target genes were normalized to that of the reference gene *RPL27* using the delta-delta-Ct method [50]. The primers used in this study are listed in the Key Resources Table.

#### RNA-seq library preparation and sequencing

Total RNA was extracted from whole cells using TRIzol Reagent following the manufacturer’s instructions. Isolated RNA was subjected to ribosomal RNA depletion using the Ribo-Zero rRNA Removal Kit (Illumina). RNA-seq libraries were prepared using the NEXTflex Rapid Directional qRNA-Seq Kit (Bioo Scientific) following the manufacturer’s instructions and sequenced using 75 bp single-end sequencing on the Illumina NextSeq 500 platform by the BioFrontiers Institute Next-Gen Sequencing Core Facility.

#### Ribosome footprinting

Ribo-seq was performed as described previously [32] using six 70% confluent 10 cm dishes of MB135-iDUX4 cells per condition. Briefly, cells were washed with ice-cold phosphate-buffered saline (PBS) supplemented with 100 μg/mL cycloheximide (Sigma-Aldrich), flash frozen on liquid nitrogen, and lysed in Lysis Buffer (PBS containing 1% (v/v) Triton X-100 and 25 U/mL TurboDNase (Ambion)). Cells were harvested by scraping and further lysed by trituration ten times through a 26-gauge needle. The lysate was clarified by centrifugation at 20,000 g for 10 min at 4 °C. The supernatants were flash frozen in liquid nitrogen and stored at −80 °C. Thawed lysates were treated with RNase I (Ambion) at 2.5 U/μL for 45 min at room temperature with gentle mixing. Further RNase activity was stopped by addition of SUPERaseIn RNase Inhibitor (Thermo Fisher Scientific). Next, ribosome complexes were enriched using MicroSpin S-400 HR Columns (GE Healthcare) and RNA extracted using the Direct-zol RNA Miniprep Kit (Zymo Research). Ribo-Zero rRNA Removal Kit was used to deplete rRNAs and the ribosome-protected fragments were recovered by running them in a 17% Urea gel, staining with SYBR Gold (Invitrogen), and extracting nucleic acids that are 27 to 30 nucleotides long from gel slices by constant agitation in 0.3 M NaCl at 4 °C overnight. The recovered nucleic acids were precipitated with isopropanol using GlycoBlue Coprecipitant (Ambion) as carrier and treated with T4 polynucleotide kinase (Thermo Fisher Scientific). Libraries were prepared using the NEXTflex Small RNA-Seq Kit v3 (Bioo Scientific) following the manufacturer’s instructions and sequenced using 75 bp single-end reads on an Illumina NextSeq 500 by the BioFrontiers Institute Next-Gen Sequencing Core Facility.

#### RNA-seq and Ribo-seq data analysis

Fastq files were stripped of the adapter sequences using cutadapt. UMI sequences were removed, and reads were collapsed to fasta format. Reads were first aligned against rRNA (accession number U13369.1), and to a collection of snoRNAs, tRNAs, and miRNA (retrieved using the UCSC table browser) using bowtie2 [51]. Remaining reads were mapped to the hg38 version of the genome (without scaffolds) using STAR 2.6.0a [52] supplied with the GENCODE 25 .gtf file. A maximum of two mismatches and mapping to a minimum of 50 positions was allowed. De-novo splice junction discovery was disabled for all datasets. Only the best alignment per each read was retained. Quality control and read counting of the Ribo-seq data was performed with Ribo-seQC [53].

Differential gene expression analysis of the RNA-seq data was conducted using DESeq2 [54]. Briefly, featureCounts from the subread R package [55] was used to assign aligned reads (in BAM format) to genomic features supplied with the GENCODE 25. gtf file. The featureCounts output was then supplied to DESeq2 and differential expression analysis was conducted with the 0 h time point serving as the reference sample. Genes with very low read count were filtered out by requiring at least a total of 10 reads across the 12 samples (3 replicates each of the 0, 4, 8, and 14 h samples). Log2 fold change shrinkage was done using the apeglm function [56].

Differential analysis of the RNA-seq and Ribo-seq data was performed using DESeq2, as previously described [57, 58], using an interaction model between the tested condition and RNA-seq - Ribo-seq counts. Only reads mapping uniquely to coding sequence regions were used. In addition, ORFquant [32] was used to derive de-novo isoform-specific translation events, by pooling the Ribo-seQC output from all Ribo-seq samples, using uniquely mapping reads. DEXSeq [33] was used to perform differential exon usage along the DUX4 time course data, using Ribo-seq counts on exonic bins and junctions belonging to different ORFquant-derived translated regions. NMD candidates were defined by ORFquant as open reading frames ending with a stop codon upstream of an exon-exon junction.

#### GO category analysis

Gene Ontology (GO) analysis was conducted using the web tool http://geneontology.org, powered by pantherdb.org. Briefly, statistical overrepresentation test using the complete GO biological process annotation dataset was conducted and p-values were calculated using the Fisher’s exact test and False Discovery Rate was calculated by the Benjamini-Hochberg procedure.

#### Polysome profiling

Polysome profiling was performed as previously described [59, 60] with the following modifications. Four 70% confluent 15 cm dishes of MB135-iDUX4 cells per condition were treated with 100 μg/mL cycloheximide for 10 min, transferred to wet ice, washed with ice-cold PBS containing 100 μg/mL cycloheximide, and then lysed in 400 μL Lysis Buffer (20 mM HEPES pH 7.4, 15 mM MgCl2, 200 mM NaCl, 1% Triton X-100, 100 μg/mL cycloheximide, 2 mM DTT, and 100 U/mL SUPERaseIn RNase Inhibitor) per 15 cm dish. The cells and buffer were scraped off the dish and centrifuged at 13,000 rpm for 10 min at 4 °C. Lysates were fractionated on a 10% to 60% sucrose gradient using the SW 41 Ti Swinging-Bucket Rotor (Beckman Coulter) at 36,000 rpm for 3 h and 10 min. Twenty-four fractions were collected using a Gradient Station ip (BioComp) and an FC 203B Fraction Collector (Gilson) with continuous monitoring of absorbance at 254 nm. RNA from each fraction was extracted using TRIzol LS Reagent (Thermo Fisher Scientific) following the manufacturer’s instructions. RT-qPCR was carried out as described above.

#### Protein extraction

Total protein was extracted from whole cells using TRIzol Reagent following the manufacturer’s instructions, excepting that protein pellets were dissolved in Protein Resuspension Buffer (0.5 M Tris base, 5% SDS). Isolated protein was quantified using the Pierce BCA Protein Assay Kit (Thermo Fisher Scientific) according to the manufacturer’s protocol. Protein was mixed with 4X NuPAGE LDS Sample Buffer (Thermo Fisher Scientific) containing 50 mM DTT and heated to 70 °C before immunoblotting.

#### Immunoprecipitation

MB135-iDUX4 myoblasts were treated with or without doxycycline for 14 h and then trypsinized prior to lysis on ice in 1 mL of Lysis Buffer (50 mM Tris-HCl pH 7.5, 150 mM NaCl, 1% NP-40) containing protease inhibitors (Sigma Aldrich). Lysates were precleared using Protein G Sepharose (Thermo Fisher Scientific) for 1 h prior to an overnight incubation at 4 °C with either anti-SRSF3 or anti-SRSF3-TR antibody. Protein G Sepharose was added the following morning for 5 h to bind the antibody, and beads were subsequently washed 5 times with 1 mL cold Lysis Buffer. After the final wash, 4X NuPAGE LDS Sample Buffer containing 50 mM DTT was added directly to the beads and samples heated to 70 °C for protein elution before immunoblotting.

#### Immunoblotting

Protein was run on NuPAGE Bis-Tris precast polyacrylamide gels (Thermo Fisher Scientific) alongside PageRuler Plus Prestained Protein Ladder (Thermo Fisher Scientific) and transferred to Odyssey nitrocellulose membrane (LI-COR Biosciences). Membranes were blocked in Intercept (PBS) Blocking Buffer (LI-COR Biosciences) before overnight incubation at 4 °C with primary antibodies diluted in Blocking Buffer containing 0.2% Tween 20. Membranes were incubated with IRDye-conjugated secondary antibodies (LI-COR Biosciences) for 1 h and fluorescent signal visualized using a Sapphire Biomolecular Imager (Azure Biosystems) and Sapphire Capture software (Azure Biosystems). When appropriate, membranes were stripped with Restore Western Blot Stripping Buffer (Thermo Fisher Scientific) before being re-probed. Band intensities were quantified by densitometry using ImageJ [61].

#### Immunofluorescence

Cells were fixed in 10% Neutral Buffered Formalin (Research Products International) for 30 min and permeabilized for 10 min in PBS with 0.1% Triton X-100. Samples were then incubated overnight at 4 °C with primary antibodies, followed by incubation with 488- or 594-conjugated secondary antibodies for 1 h prior to counterstaining and mounting with Prolong Diamond Antifade Mountant with DAPI (Thermo Fisher Scientific). Slides were imaged with a DeltaVision Elite deconvolution microscope, CoolSNAP HQ^2^ high-resolution CCD camera, and Resolve3D softWoRx-Acquire v7.0 software. Image J software [61] was used for image analysis.

#### Solid phase synthesis of TMO chimeras (ASOs)

TMO chimeras were synthesized according to the previously reported procedure [39, 40]. Briefly, the 5’-dimethoxytrityl (DMT) protecting group of the solid supported 2’-deoxyribonucleoside (CPG-500 support, Glen Research) was deprotected in the first stage by using 3% trichloroacetic acid in dichloromethane. In the second stage, condensation of the resulting CPG-500 support linked 5’-hydroxyl-2’-deoxyribonucleoside with the 6’-DMT-morpholinonucleoside 3’-phosphordiamidites of mA^Bz^, mG^iBu^, mC^Bz^, mT (ChemGenes) or commercial 2’-deoxyribonucleoside 3’-phosphoramidites was achieved using 5-ethylthio-1H-tetrazole (ETT) in anhydrous acetonitrile as activator (30 sec condensation time). Subsequent conversion of P(III) linkages to P(V) thiophosphoramidate (TMO) or P(V) 2’-deoxyribonucleoside 3’-thiophosphate was achieved by using 3-[(Dimethylaminomethylene)amino]-3H-1,2,4-dithiazole-5-thione (DDTT) as the sulfurization agent. Finally, the unreacted hydroxyl groups were acetylated by conventional capping reagents (Cap A: Tetrahydrofuran/Acetic Anhydride and Cap B: 16% 1-Methylimidazole in Tetrahydrofuran; Glen Research). The 5’-DMT protecting group on the resulting dinucleotide was next deprotected using deblocking mixture and this DMT deprotected dinucleotide was then used for additional cycles in order to generate ASOs having internucleotide thiophosphoramidate or thiophosphate linkages. The above cycle was repeated to provide the thiomorpholino oligonucleotide chimeras of the desired length and sequence. Cleavage of these 5’-protected DMT-on oligonucleotides from the solid support and deprotection of base and phosphorus protecting groups was carried out using 0.5 mL of 28% aqueous ammonia at 55 °C for 16 h. Subsequently, the CPG was filtered through a micro spin centrifuge filter with pore size of 0.2 μm and the resulting filtrate was evaporated to dryness on a SpeedVac (Thermo Fisher Scientific). The residue was dissolved in 0.75 mL of 3% acetonitrile/water mixture and filtered through a micro spin centrifuge filter. A small portion of the crude sample was withdrawn and submitted to LCMS analysis. The remaining reaction mixture was purified by RP-HPLC. Fractions containing the pure ASO were combined, evaporated to dryness, and submitted to LCMS analysis. Fractions containing the pure DMT-on ASO were dissolved in 0.5 mL of detritylation mixture. After 25 min at 40 °C, the mixture was neutralized with 5 μL of triethylamine, filtered using a micro spin centrifuge filter, and the filtrate containing the sample was purified by RP-HPLC column chromatography. Fractions containing the final DMT-off product were combined and evaporated to dryness on a SpeedVac. The residue was submitted to LCMS analysis in order to determine the purity of the sample. The concentration of the ASO was determined by NanoDrop spectrophotometry before storing the samples at −20 °C.

#### LCMS analysis

LCMS analysis was performed on an Agilent 6530 series Q-TOF LC/MS spectrometer. A Waters ACQUITY UPLC® BEH C18, 1.7 μm, 2.1 X 100 nm column was used as the stationary phase. Aqueous phase was Buffer A (950 mL water, 25 mL methanol, 26 mL hexafluoro-2-propanol (HFIP) and 2.5 mL triethyl amine) and organic phase was Buffer B (925 mL methanol, 50 mL water, 26 mL hexafluoro-2-propanol (HFIP) and 2.5 mL triethyl amine). The gradient was 0-100% of Buffer B for 30 min followed by 100% Buffer B for 5 min at a flow rate of 0.2 mL/min and a set temperature of 25 °C. The observed masses of the ASOs were consistent with the expected theoretical masses.

#### Antisense oligonucleotide transfections

ASOs were transfected into MB135-iDUX4 cells 40 h prior to doxycycline induction using Lipofectamine RNAiMAX Transfection Reagent (Thermo Fisher Scientific) following the manufacturer’s instructions. The ASOs used in this study are listed in the Key Resources Table.

#### Cell viability assays

Trypan blue dye was used to determine the viability of MB135-iFLAG-SRSF3-FL and -TR cell lines. Ten microliters of trypsinized and resuspended cells were mixed with 10 μL of 0.4% Trypan Blue Stain (Thermo Fisher Scientific) for 1 min before immediate counting using a hemocytometer and Motic AE2000 inverted light microscope. Caspase 3/7 activity was used to determine the viability of MB135-iDUX4 myoblasts treated with ASOs or proteasome inhibitors. MB135-iDUX4 cells were seeded in 24-well plates at 8e4 cells per well, transfected with ASOs as described above, and 40 h later treated with 2 μg/mL doxycycline hyclate or seeded in 96-well plates at 3e3 cells per well and 24 h later treated with 1 μg/mL doxycycline hyclate and either DMSO or 10 μM MG132 (Sigma-Aldrich). Caspase 3/7 activity was measured 16 and 24 h later using the Caspase-Glo 3/7 Assay System (Promega) following the manufacturer’s instructions. Luminescence was detected using a GloMax-Multi Detection System (Promega).

#### Antibodies

The antibodies used in this study are anti-DUX4 (Abcam 124699), anti-Histone H3 (Abcam 1791), anti-SRSF3 (Thermo Fisher Scientific 33-4200), anti-SRSF3-TR (this paper), anti-RENT1/hUPF1 (Abcam ab109363), Drop-n-Stain CF 488A Donkey Anti-Rabbit IgG (Biotium 20950), Drop-n-Stain CF 594 Donkey Anti-Rabbit IgG (Biotium 20951), IRDye 650 Goat anti-Mouse IgG Secondary Antibody (LI-COR Biosciences 926-65010), and IRDye 800CW Goat anti-Rabbit IgG Secondary Antibody (LI-COR Biosciences 926-32211).

### QUANTIFICATION AND STATISTICAL ANALYSIS

#### Data analysis, statistical tests, and visualization

All data analysis and statistical tests were performed in the R programming environment and relied on Bioconductor [62] and ggplot2 [63]. Plots were generated using R plotting functions and/or the ggplot2 package. Bar graphs were generated using GraphPad Prism software version 9.0. Biological replicates were defined as experiments performed separately on distinct samples (i.e. cells cultured in different wells) representing identical conditions and/or time points. No outliers were eliminated in this study. Statistical tests were performed using R functions or GraphPad Prism.

## ACKNOWLEDGEMENTS

We thank Stephen Tapscott for the MB135-iDUX4/ZSCAN4-mCherry cell line. We thank Jeffrey Kieft for his guidance in carrying out polysome profiling. We thank Neelanjan Mukherjee, Olivia Rissland, Srinivas Ramachandran, and all members of the Jagannathan laboratory for insightful manuscript feedback. We thank the BioFrontiers Institute Next-Gen Sequencing Core Facility, which performed the Illumina sequencing and library construction. This work was supported by the RNA Bioscience Initiative, University of Colorado Anschutz Medical Campus (S.J.), the University of Colorado Boulder (K.S. and M.H.C.), Friends of FSH Research and The Chris Carrino Foundation for FSHD AWD-194864 (S.J.), the National Institutes of Health grant R35GM133433 (S.J.), the FSHD Society FSHS-82018-01 (A.E.C. and M.D.), the California Tobacco-Related Disease Research Grants Program 27KT-0003 (S.N.F.), and the National Institutes of Health DP2GM132932 (S.N.F.).

## AUTHOR CONTRIBUTIONS

A.E.C., M.C.D., and S.J. conceived and designed the study. A.E.C, M.C.D., M.A.C., and T.F. performed experiments. K.S. and M.H.C. provided the thiomorpholino oligonucleotides. A.E.C., L.C., T.M., R.F., A.E.G., M.H.C., S.N.F., and S.J. analyzed data. A.E.C. and S.J. wrote the paper with input from all authors.

## DECLARATION OF INTERESTS

The authors declare no competing interests.

## SUPPLEMENTAL FILES

**Figure S1.**
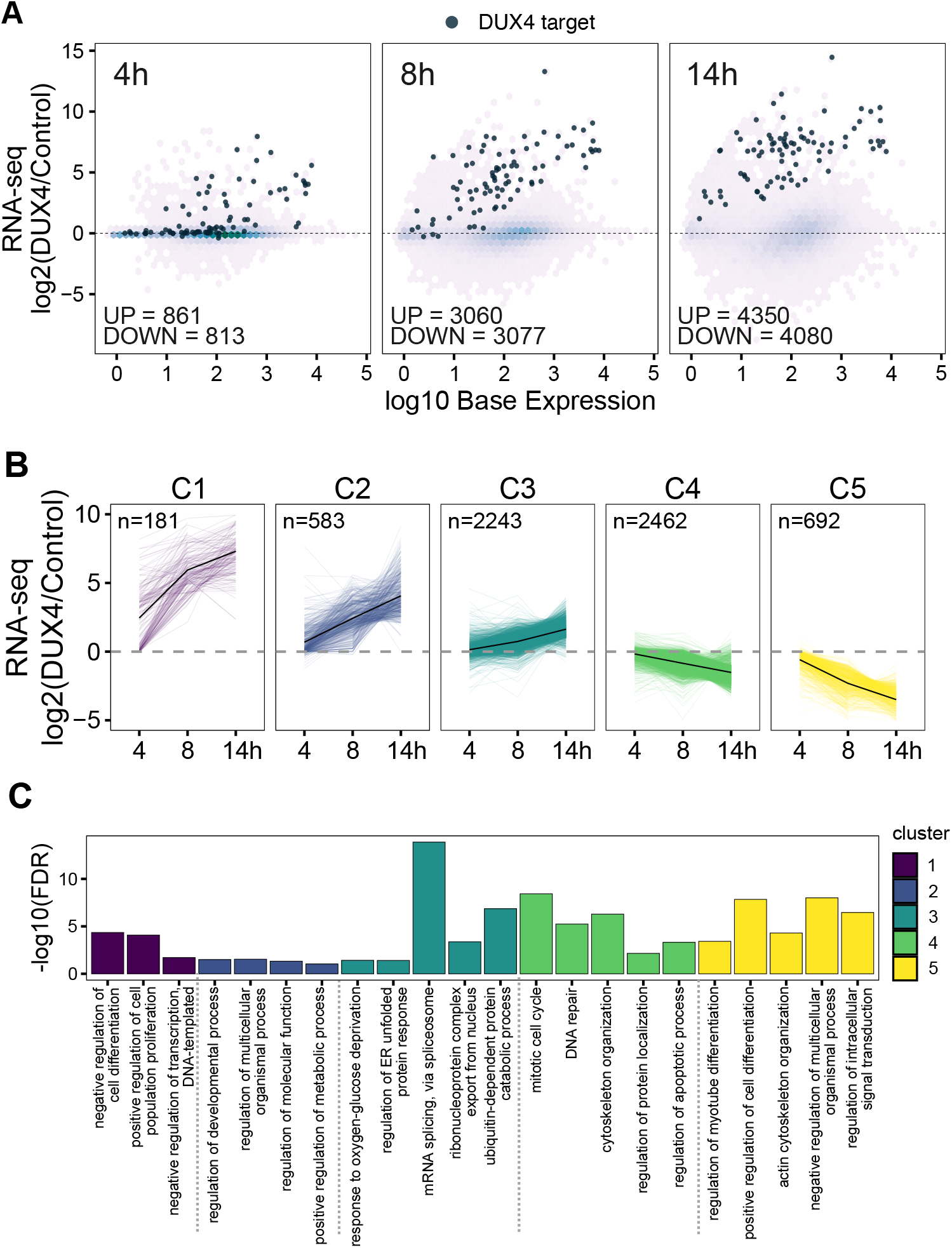
Time course RNA-seq in MB135-iDUX4 myoblasts reveals early transcript-level changes in pathways underlying FSHD pathology. (**A**) M-A plots for RNA-seq data after 4, 8, and 14 h of DUX4 induction compared to the 0 h control. DUX4 target status defined as in [25]. (**B**) Log2 fold change in RNA expression from the 0 h time point is shown for each gene after k-means clustering. The thick black line represents the cluster mean. (**C**) GO analysis results of selected gene sets (biological process complete) that are significantly enriched in each cluster defined in (**B**).

**Figure S2.**
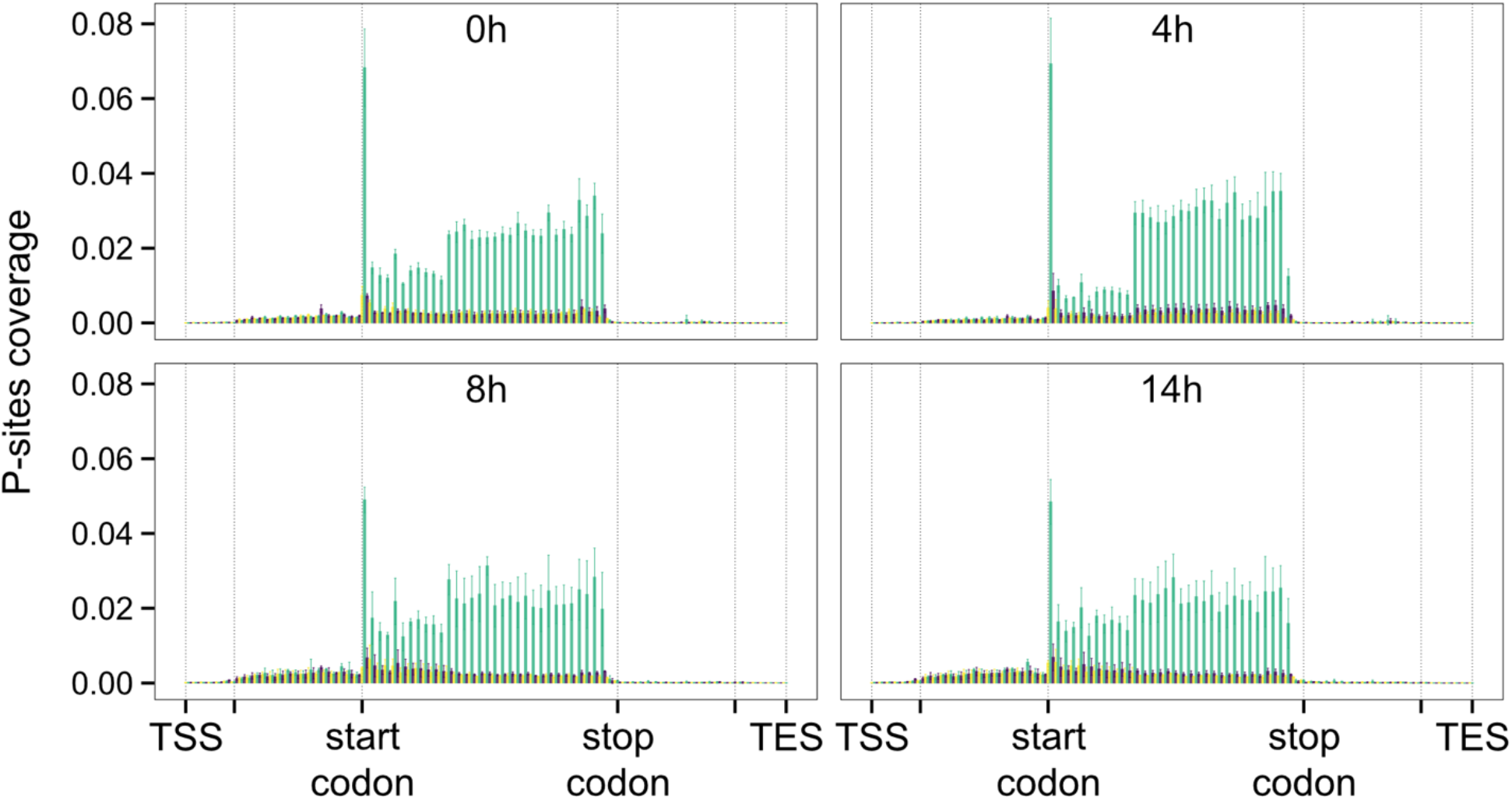
Ribo-seq quality control. Aggregate profile of P-sites coverage (as calculated by Ribo-seQC) depicting single nucleotide resolution of Ribo-seq data along the time course. Each frame is shown with a different color. Error bars represent the standard deviation from three biological replicates.

**Figure S3.**
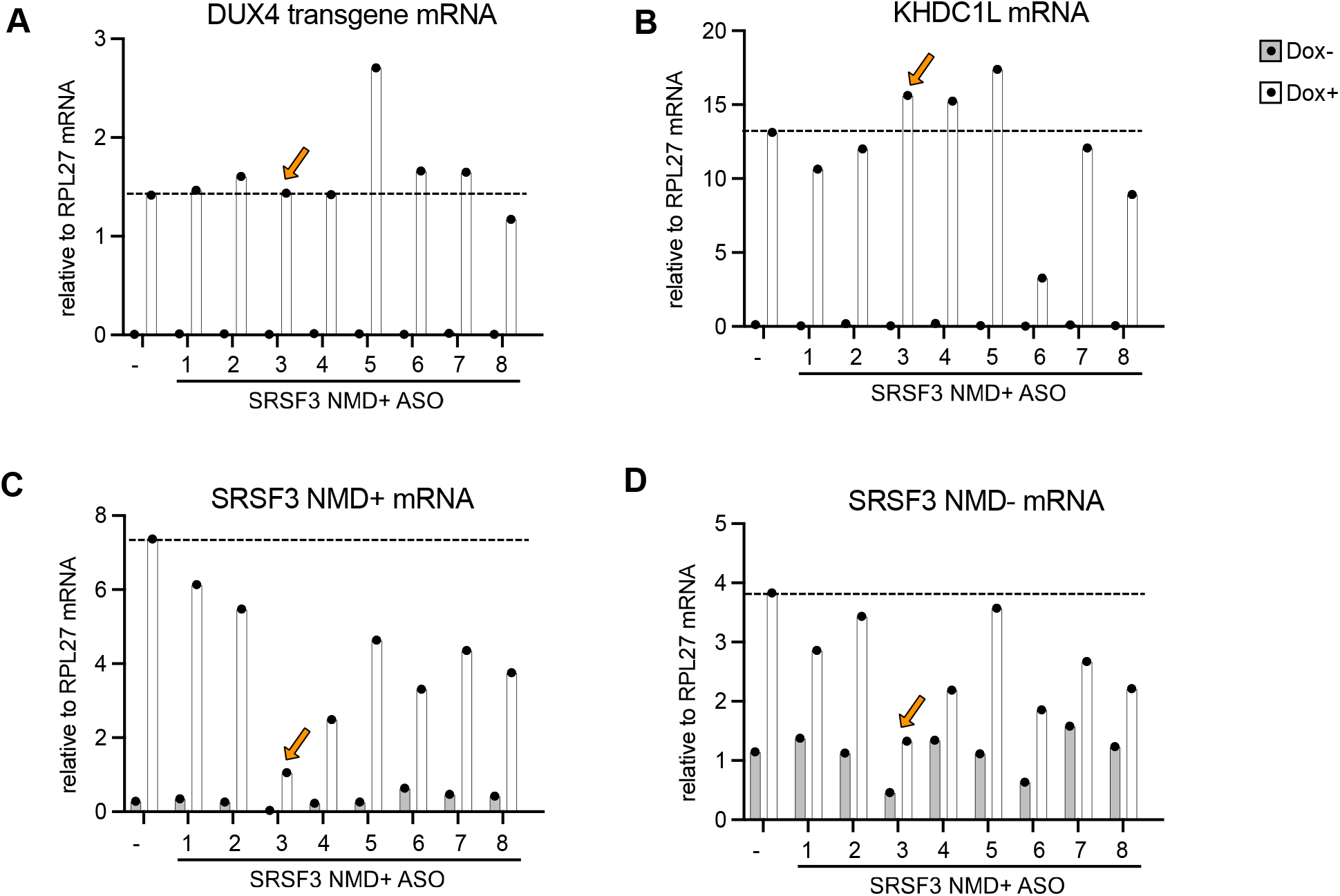
Relative RNA levels of (**A**) DUX4 transgene, (**B**) KHDC1L (DUX4 target gene), (**C**) SRSF3 NMD+, and (**D**) SRSF3 NMD-isoforms as determined by RT-qPCR following transfection with antisense oligos (ASOs) targeting SRSF3 NMD+ and doxycycline (Dox) treatment for 14 h to induce DUX4 in MB135-iDUX4 myoblasts. The tested ASOs are numbered 1-8; “-“ indicates no ASO control. The orange arrow indicates the ASO chosen for further studies in Figure 4F.

**Video 1. Time course imaging following DUX4 expression.**

Live cell fluorescence microscopy recording of MB135-iDUX4/ZSCAN4-mCherry myoblasts treated with doxycycline to induce DUX4 expression.

**Supplementary Table 1.**

DESeq2 differential gene expression analysis results for DUX4 time course RNA-seq data at 4, 8, or 14 h post-induction compared to 0 h (control).

**Supplementary Table 2.**

Cluster analysis of RNA-seq log2 fold change at 4, 8, or 14 h of DUX4 induction compared to 0 h (control); and Gene Ontology analysis of genes within each cluster.

**Supplementary Table 3.**

ORFquant analysis results for DUX4 time course RNA-seq and Ribo-seq data at 4, 8, or 14 h post-induction compared to 0 h (control); and Gene Ontology analysis of genes with downregulated ribosome density at 14 h.

**Supplementary Table 4.**

DEXSeq analysis results for DUX4 time course Ribo-seq data at 4, 8, or 14 h post-induction compared to 0 h (control); and Gene Ontology analysis of NMD targets upregulated at 14 h.

## APPENDIX

### KEY RESOURCES TABLE

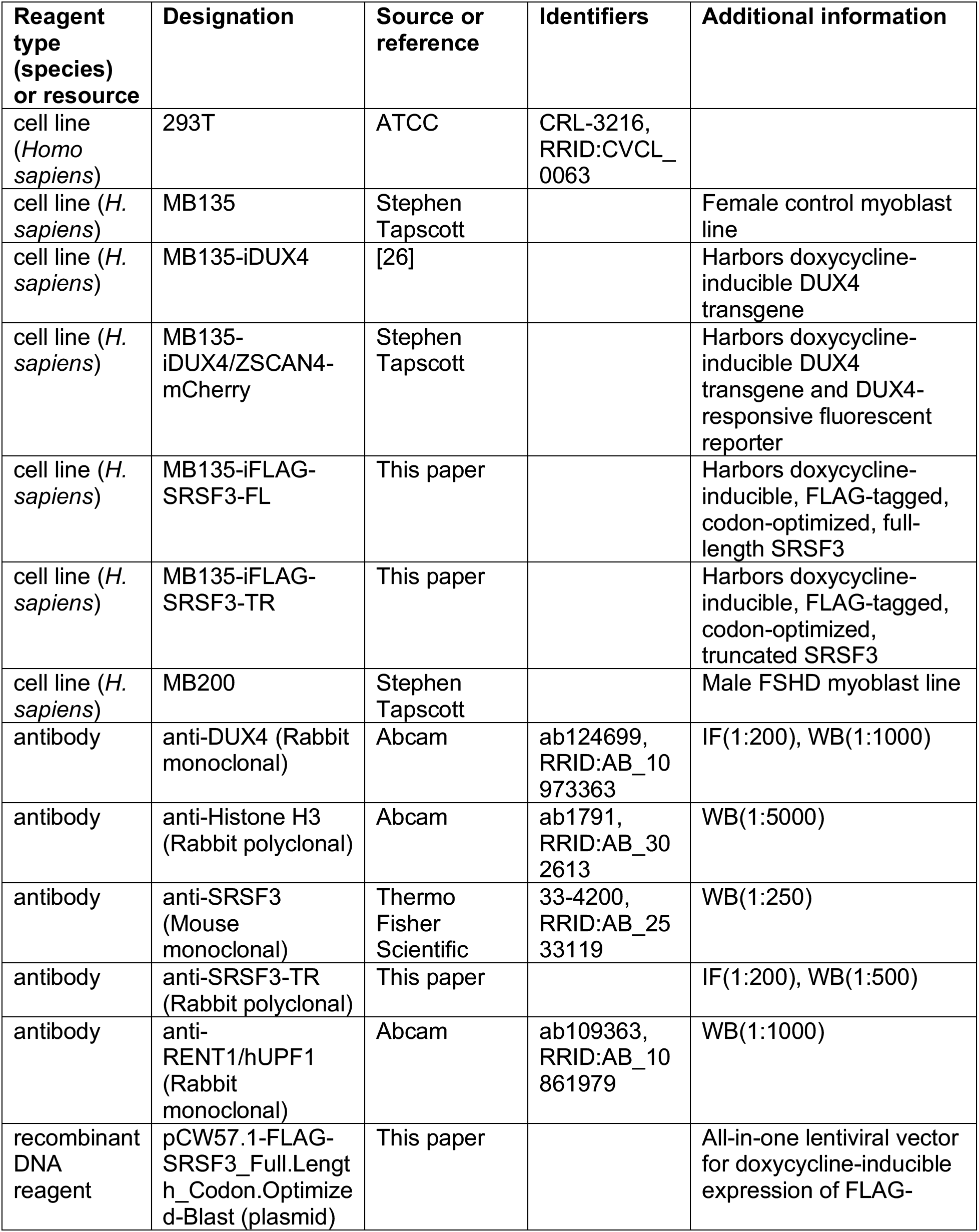
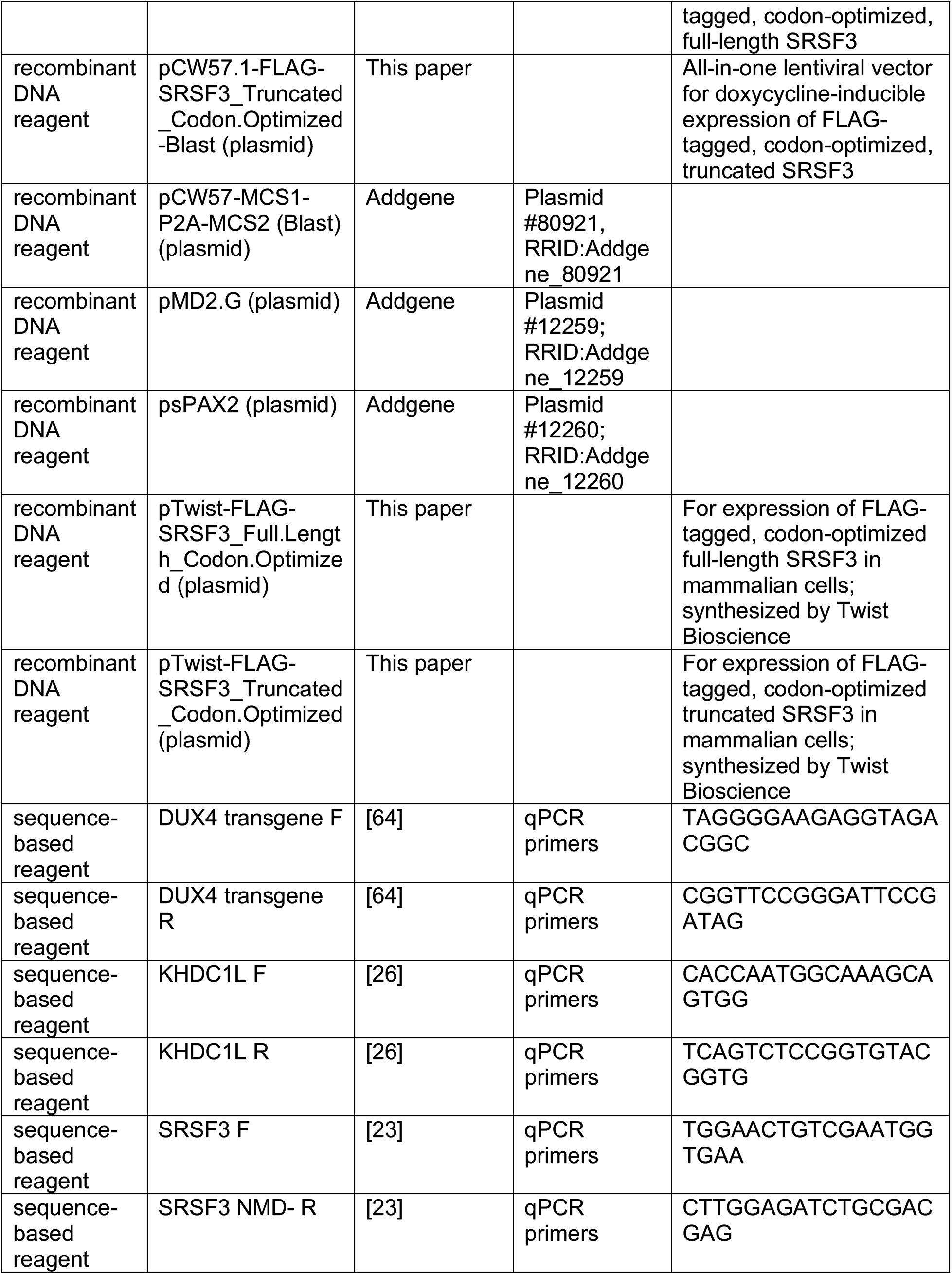
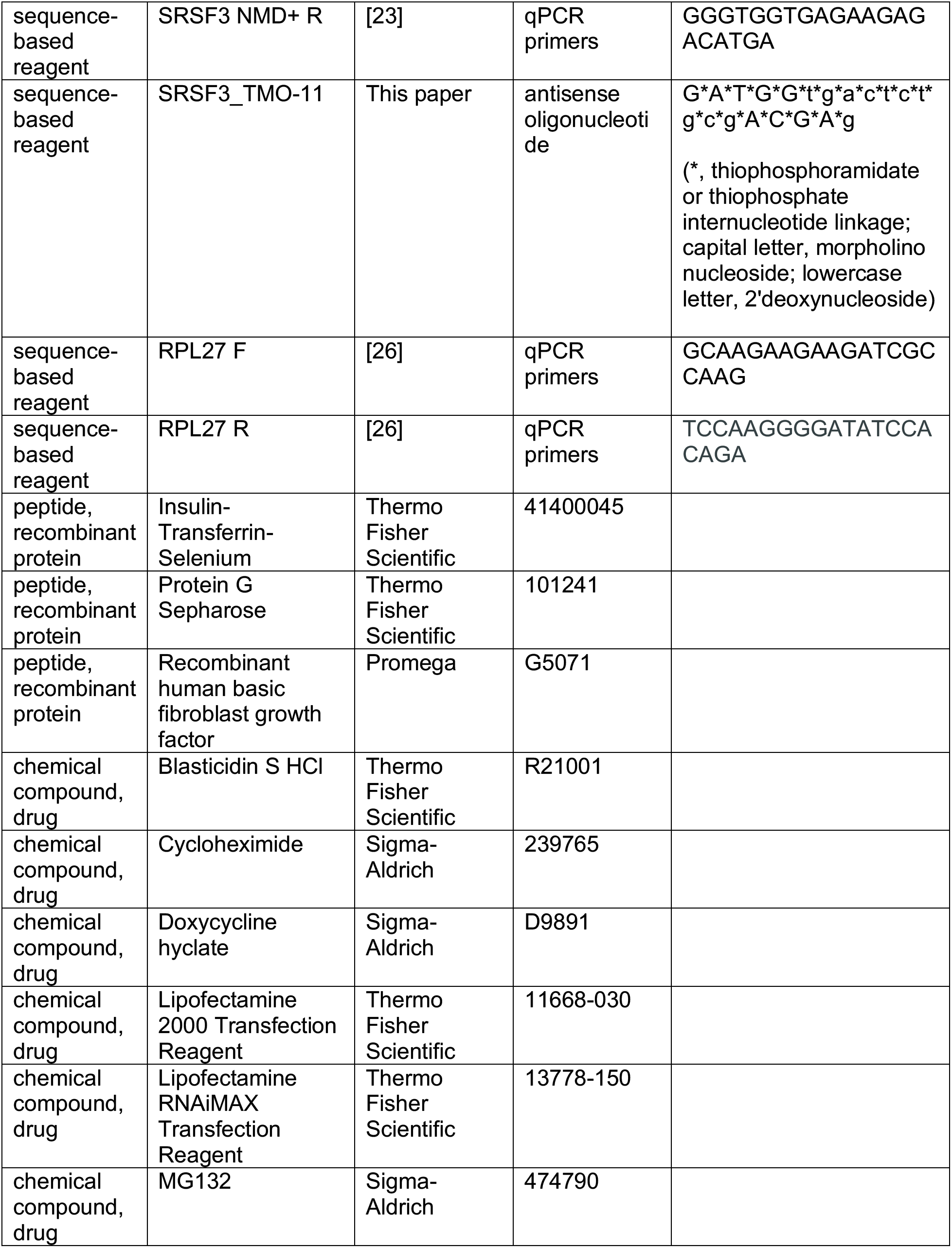
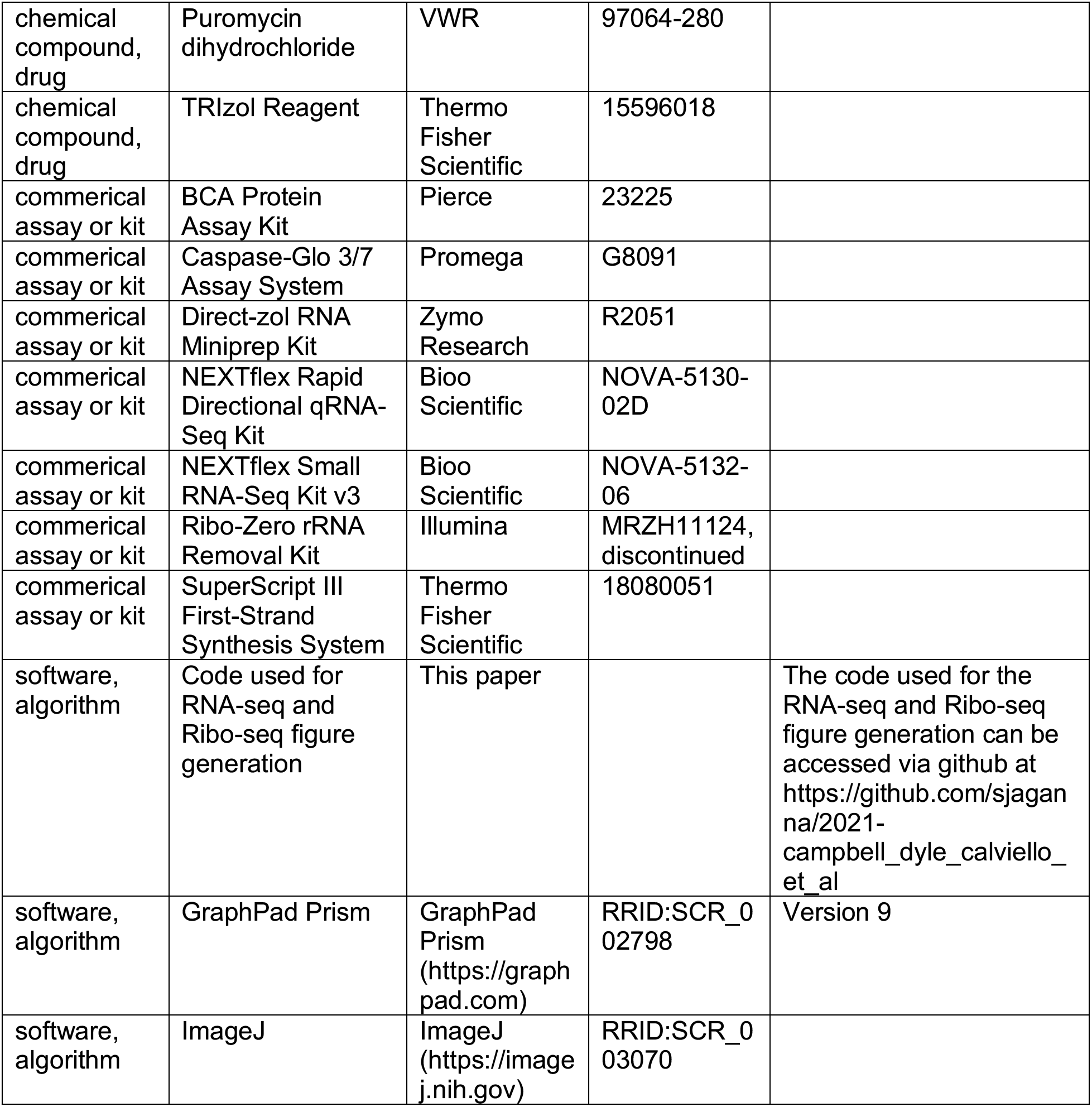

## Notes

### Competing Interest Statement

The authors have declared no competing interest.

### Summary of Updates

Substantial revision to manuscript framing and additional data.

